# *Rhizobium moroccans* sp. nov.: a plant-growth-promoting endophyte from *Peganum harmala* in an arid environment

**DOI:** 10.64898/2026.02.23.707546

**Authors:** Salma Mouhib, Khadija Ait Si Mhand, Juan Carlos Fernández-Cadena, Mohamed Hijri

## Abstract

The diversity and ecological roles of *Rhizobium* species beyond legume symbiosis remain insufficiently characterized, particularly in arid ecosystems. Here, we report the isolation and comprehensive characterization of strain AGC32, an endophytic bacterium recovered from surface-sterilized roots of the medicinal xerophyte *Peganum harmala* growing in Moroccan drylands. The isolate is a non-halophilic, aerobic, Gram-negative bacterium with optimal growth at 30–35°C, pH 5.0, and 1% NaCl. Phylogenomic analysis placed AGC32 within the genus *Rhizobium*. However, average nucleotide identity values below 96%, digital DNA–DNA hybridization values below 70%, and the absence of assignment to any validly published type strain supported its classification as a novel species, for which the name *Rhizobium moroccans* sp. nov. is proposed. Comparative genomics revealed substantial genome rearrangements relative to its closest relative, *Rhizobium deserti*, indicating distinct evolutionary trajectories. The high-quality draft genome encodes pathways associated with nitrogen limitation (including complete allantoin utilization), polyphosphate metabolism, oxidative and osmotic stress tolerance, and organic acid utilization. Phenotypic assays corroborated genomic predictions, demonstrating preferential metabolism of organic acids, utilization of carbohydrates, and plant growth–promoting traits including nitrogen fixation and solubilization of phosphorus, potassium, and silicon. These findings expand the ecological knowledge of *Rhizobium* and reveal adaptation to a non-legume host in an arid environment.

**IMPORTANCE:** Bacteria of the genus *Rhizobium* are widely known for forming nitrogen-fixing symbioses with legumes. However, their roles in non-leguminous desert plants remain poorly understood. We isolated and characterized a new species, *Rhizobium moroccans*, from the roots of the medicinal plant *Peganum harmala* growing in Moroccan drylands. This bacterium shows genetic and physiological traits that support survival under drought and nutrient limitation and displays multiple plant-beneficial properties. Our results demonstrate that *Rhizobium* species are not restricted to legumes but can form intimate associations with medicinal plants in extreme environments. The discovery of this novel species highlights desert plants as reservoirs of previously unrecognized microbial diversity and suggests potential applications for sustainable agriculture in arid regions.

## INTRODUCTION

Plant roots harbor highly structured microbial communities that play critical roles in nutrient acquisition, stress tolerance, and plant fitness. Within these communities, the root endosphere represents a strongly filtered niche enriched in bacterial taxa capable of intimate host colonization (1, 2). In arid and semi-arid ecosystems, this filtering is intensified by environmental stressors such as drought, salinity, and nutrient limitation, which shape both microbial diversity and functional traits (3, 4). Therefore, desert-adapted plants constitute promising reservoirs of stress-tolerant and metabolically versatile microorganisms (5).

Among root-associated bacteria, the genus *Rhizobium* is best known for nitrogen-fixing symbioses with legumes. However, accumulating genomic and ecological evidence indicates that *Rhizobium* species extend far beyond classical nodulating lifestyles (2). Their multipartite genomes, comprising a conserved chromosome and accessory plasmids or chromids, facilitate horizontal gene transfer and ecological diversification (6, 7). Evolutionary analyses suggest that ancestral Rhizobiales were generalist root colonizers that later acquired nodulation genes, supporting repeated transitions between free-living, endophytic, and symbiotic states (8).

Consistent with this evolutionary plasticity, *Rhizobium* lineages are frequently detected in non-legume roots, including plants adapted to extreme environments. In Moroccan drylands, the root endosphere of *Malva sylvestris* L. was reported to be dominated by *Rhizobium* despite the absence of nodulation (2). Similar enrichment of Rhizobiales has been documented in desert plants, where endophytic bacteria contribute to plant growth promotion and stress tolerance (3, 9). In non-legume hosts, rhizobia typically express plant growth–promoting traits, including phytohormone production, phosphate solubilization, siderophore secretion, and ACC deaminase activity, rather than symbiotic nodulation pathways (10-12). These observations indicate that rhizobia occupy broader ecological niches than traditionally recognized.

Despite growing recognition of rhizobial ecological flexibility, the diversity and genomic basis of *Rhizobium* adaptation to non-legume medicinal plants in arid environments remain poorly characterized. Medicinal xerophytes are particularly relevant because their production of bioactive secondary metabolites may impose additional selective pressures on associated microbiota. *Peganum harmala* L., a perennial plant native to North African and Middle Eastern drylands, thrives in alkaline and drought-prone soils and synthesizes β-carboline alkaloids with documented antimicrobial activity (13). While its phytochemistry is well studied, its endophytic bacterial diversity, especially rhizobial lineages, has not been taxonomically or genomically resolved.

We hypothesized that the root endosphere of *P. harmala* in Moroccan drylands harbors phylogenetically distinct and genomically adapted *Rhizobium* lineages reflecting both host filtering and extreme environmental pressures. Specifically, we expected that such isolates would (i) represent novel taxonomic entities within the genus and (ii) display genomic signatures associated with stress tolerance, metabolic versatility, and plant-associated lifestyles independent of nodulation.

To test this hypothesis, we isolated an endophytic *Rhizobium* strain from *P. harmala* roots and applied a polyphasic taxonomic framework integrating whole-genome sequencing, phylogenomic reconstruction, average nucleotide identity (ANI), digital DNA–DNA hybridization (dDDH), comparative genomic analysis, functional annotation, and phenotypic characterization. Through this integrative approach, we aimed to determine its taxonomic position and elucidate genomic features underlying adaptation to a non-legume host in an arid ecosystem.

Our findings support the designation of a novel species, *Rhizobium moroccans* sp. nov. AGC32, and expand the ecological and evolutionary understanding of rhizobia beyond classical legume symbiosis. This work highlights medicinal desert plants as reservoirs of previously unrecognized rhizobial diversity and provides insight into bacterial adaptation at the soil–root interface in extreme environments.

## RESULTS

### Isolation of endophytic bacteria

Surface-sterilized root tissues of *Peganum harmala* yielded several bacterial isolates on TSA (×0.1 and ×1) and a few on PDA (×0.1 and ×1). Among these, strain AGC32 was recovered from root fragments plated on PDA ×0.1, although this medium is typically used for fungal isolation. Colonies appeared after 48–72 h of incubation at 28 °C. They were circular, smooth, convex, opaque, creamy-white, and approximately 1 mm in diameter after 72 h (Fig. S1). Cells were Gram-negative, motile coccobacilli, as observed by light microscopy, and no spore formation was detected.

### Genome-based taxonomic characterization

#### Phylogenomics and genome structure

Phylogenomic analysis based on 13 complete *Rhizobium* genomes placed strain AGC32 within the *Rhizobium* clade and clearly separated from the outgroup *Mesorhizobium jarvisii* (Fig. 1). AGC32 formed a distinct monophyletic lineage, separate from the closest type strains included in the analysis, including *Rhizobium deserti, Rhizobium puerariae, Rhizobium leguminosarum* bv. *viciae, Rhizobium hidalgonense, Rhizobium mayense*, and *Rhizobium aegyptiacum*.

**Figure 1.**
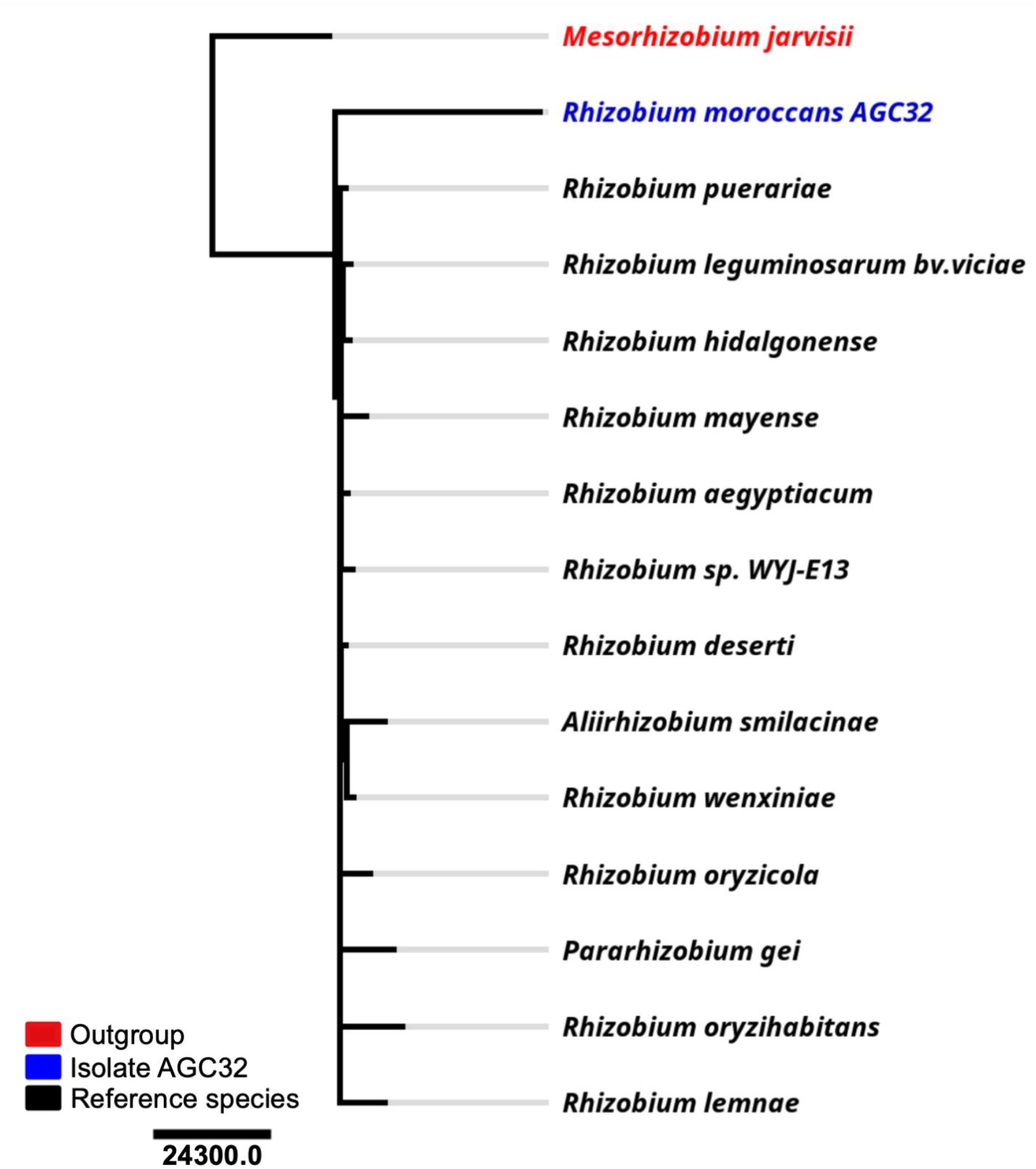
Rooted phylogenomic tree of the genus *Rhizobium* based on 13 complete genomes. Maximum-likelihood analysis positions isolate AGC32 (candidate *Rhizobium moroccans* sp. nov.) as a distinct lineage closely related to its nearest species. Colors indicate taxonomic grouping: red, outgroup (*Mesorhizobium jarvisii*); blue, the newly described species AGC32; black, reference species. The scale bar represents the number of substitutions per 100,000 nucleotide sites (scale = 24,300).

Pairwise genome comparisons showed an average nucleotide identity (ANI) of 79.9% and a digital DNA–DNA hybridization (dDDH) value of 20% relative to its closest phylogenetic neighbor (Table S1).

Whole-genome alignment using progressiveMauve revealed partial conservation of the genomic backbone relative to *R. deserti*, with 16 locally collinear blocks (LCBs) (Fig. 2).

**Figure 2.**
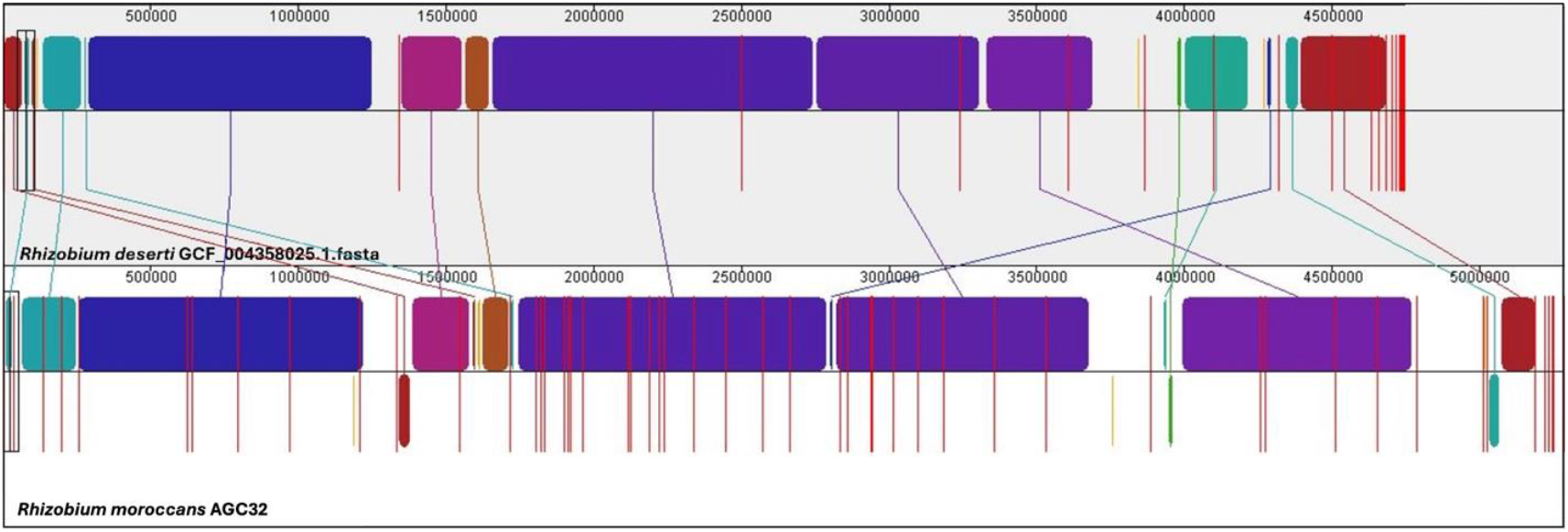
Comparative genomic synteny between *Rhizobium moroccans* AGC32 and *R. deserti*. Whole-genome alignment was performed using Mauve, highlighting significant genome rearrangements across 16 locally collinear blocks (LCBs; minimum weight = 13,100). Each colored block represents an LCB, corresponding to a homologous region conserved between the two genomes. Blocks of the same color indicate positional homology, while their vertical placement (above or below the central axis) reflects orientation relative to the reference genome (forward or inverted, respectively). Lines connecting blocks indicate conserved sequences between the genomes, illustrating inversions, translocations, and other structural variations.

Several LCBs were inverted or repositioned. Increased structural variability was observed in terminal genomic regions.

#### Genome assembly and quality metrics

The draft genome of strain AGC32 consists of 5.24 Mb distributed across 55 contigs, with an N50 of 135,068 bp and a GC content of 61.62% (Table S2). Genome completeness was estimated at 98.82% with 0.92% contamination and an average sequencing coverage of 73×.

The partial16S rRNA gene sequence showed 97.6% similarity to its closest validly named relative. Multilocus sequence typing (MLST) analysis did not match any existing allele profiles, and TYGS analysis did not affiliate AGC32 with any recognized type strain.

### Functional and metabolic potential

#### Functional Annotation

Genome annotation identified genes distributed across major metabolic categories (Fig. 3a and Table S3). The largest functional groups included protein biosynthesis (132 genes), carbohydrate metabolism (80 genes), flagellar motility (47 genes), and DNA repair (44 genes). Genes associated with oxidative stress, osmotic stress (16 genes), detoxification, and resistance to antibiotics and toxic compounds (16 genes) were also identified.

**Figure 3.**
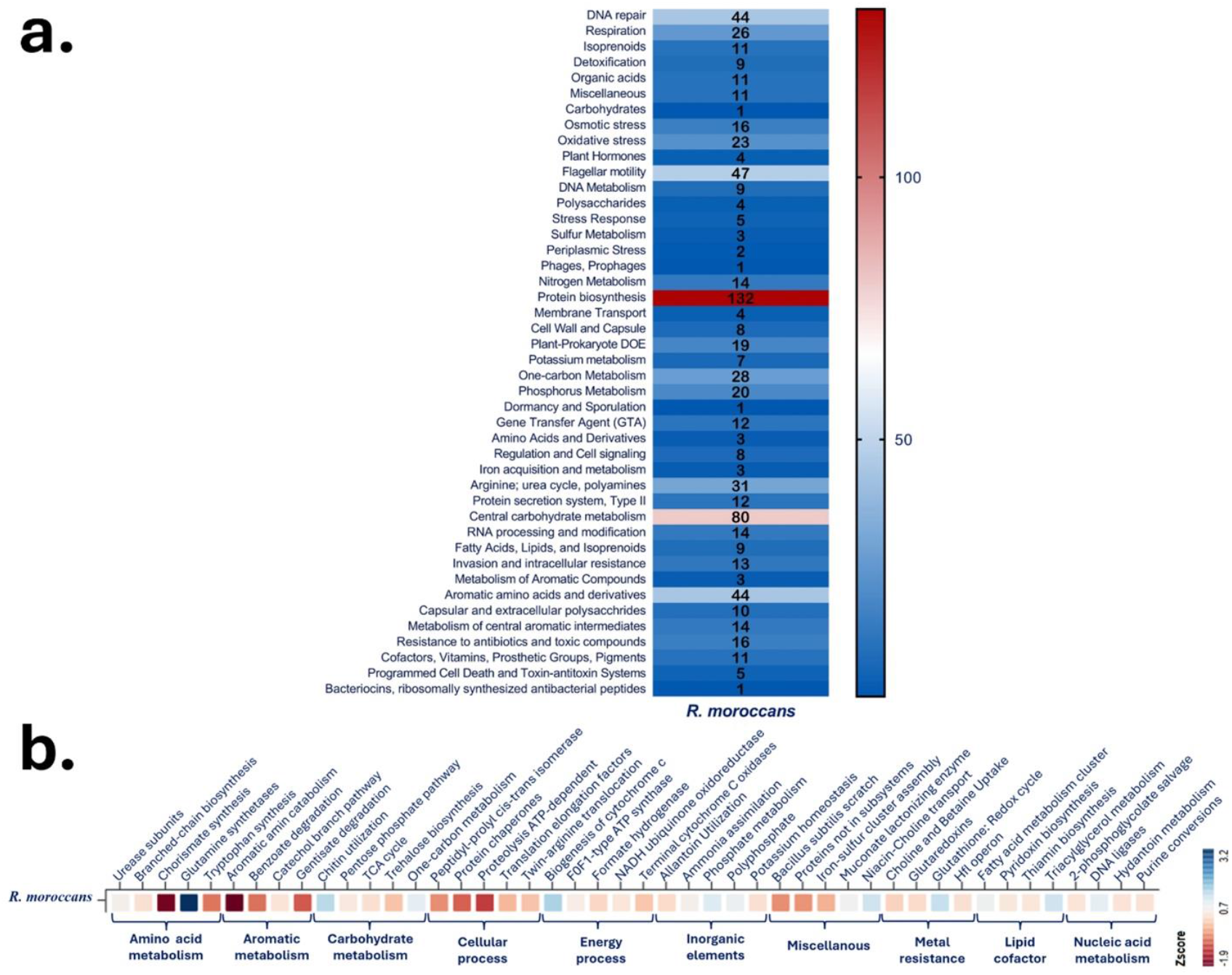
Functional and metabolic architecture of *R. moroccans* sp. nov. AGC32. (a) Distribution of functional capabilities based on KEGG ortholog (KO) gene counts grouped into major metabolic categories. (b) Z-score heatmap showing enrichment patterns across key metabolic subsystems, including amino acid metabolism, aromatic compound metabolism, carbohydrate metabolism, cellular processes, energy metabolism, inorganic ion metabolism, miscellaneous functions, metal resistance, lipid and cofactor metabolism, and nucleic acid metabolism.

Subsystem annotation revealed genes involved in nitrogen metabolism (14 genes), iron acquisition, sulfur metabolism, and inorganic ion transport.

Heatmap analysis of normalized gene counts showed representation across amino acid metabolism, carbohydrate utilization, energy production, inorganic element metabolism, lipid metabolism, metal resistance, and nucleic acid metabolism (Fig. 3b).

The genome encodes a complete denitrification pathway, high-affinity iron uptake systems (EfeUOB), glutamine synthetase variants, cytochrome c biogenesis proteins, glutathione redox components, and polyphosphate metabolism genes.

#### Secondary Metabolite Biosynthetic Gene Clusters

Genome mining using antiSMASH identified five putative secondary metabolite biosynthetic gene clusters (BGCs) distributed across the AGC32 genome (Fig. 4). These included a terpene precursor cluster (Region 1.1), a RiPP-like cluster (Region 13.1), a homoserine lactone cluster (Region 25.1), a terpene cluster (Region 36.1), and an NRPS-like/Type I PKS hybrid cluster (Region 45.1). These BGC classes are commonly associated with membrane stabilization, oxidative stress protection, antimicrobial compound production, quorum sensing (AHL synthesis), and volatile organic compound biosynthesis in rhizobial species.

**Figure 4.**
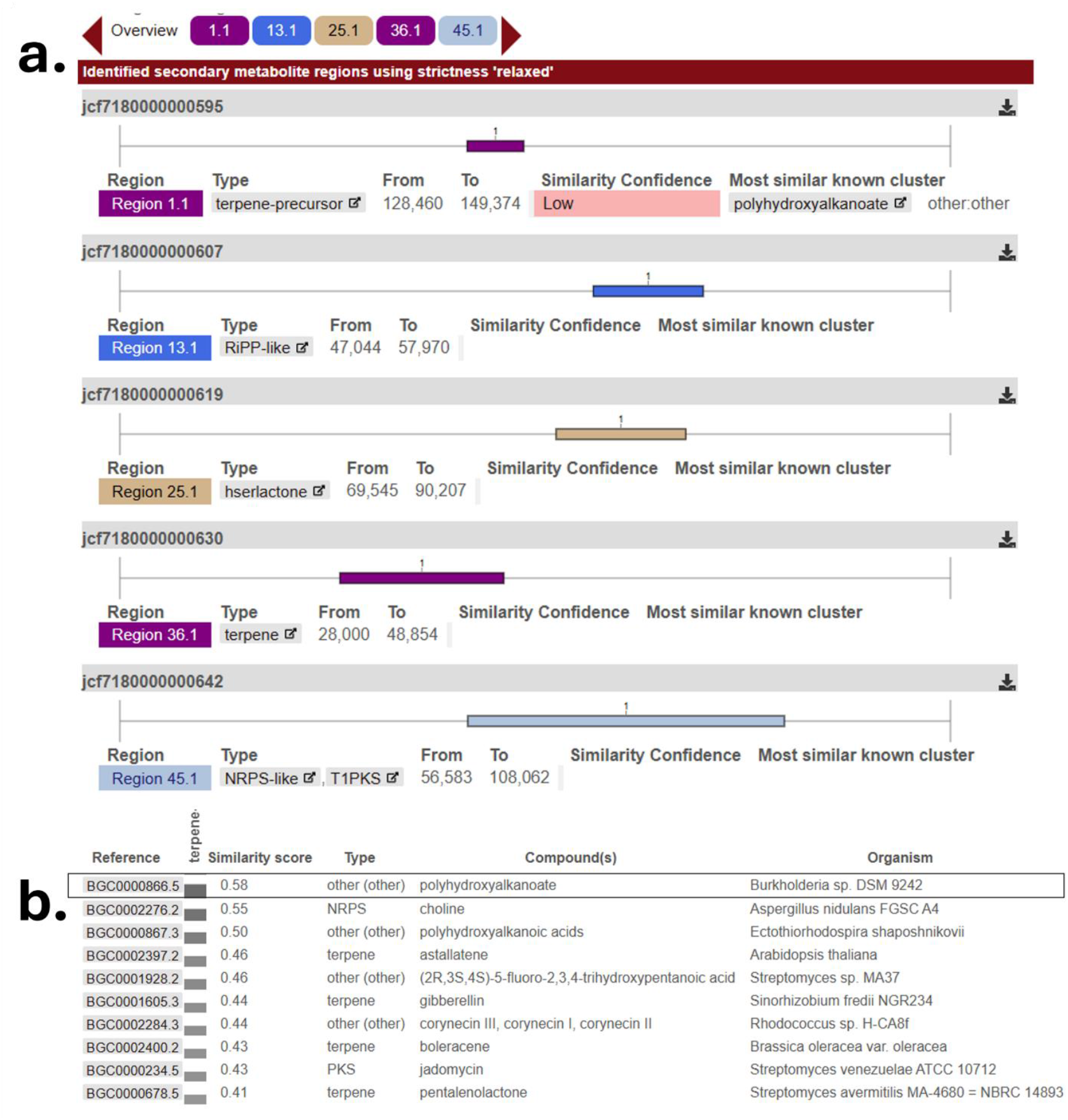
Genome mining and comparative analysis of biosynthetic gene clusters (BGCs) predicted using antiSMASH. (a) BGC regions identified in *R. moroccans* sp. nov. AGC32. (b) Top similarity scores of the predicted clusters compared to characterized BGCs in reference databases.

Similarity comparisons against the MIBiG database revealed generally low similarity scores (<0.6) to characterized clusters (Fig. 4), with the highest similarity (0.58) observed for the polyhydroxyalkanoate-associated biosynthetic region. The limited similarity to known BGCs suggests potential novelty of secondary metabolite pathways in AGC32. Overall, the genome architecture of *Rhizobium moroccans* AGC32 is consistent with a free-living, rhizosphere-associated lifestyle typical of members of the genus *Rhizobium*.

### Species description

#### *Rhizobium moroccans* sp. nov

*Rhizobium moroccans* AGC32 was isolated from the roots of *Peganum harmala*. Growth occurred on potato dextrose agar (PDA, ×0.1) at 28–35 °C within 72 h. Colonies were small (∼1 mm in diameter), circular, raised, smooth, opaque, creamy-white, mucoid, and had entire margins (Fig. S1). Cells observed under light microscopy were Gram-negative and motile coccobacilli, and no spores were detected. All phenotypic assays were performed at ∼28–30 °C under aerobic conditions.

- Etymology: M.L. neut. adj. *moroccans*; “Of Morocco,” referring to the country from which it was isolated.
- Species name: *Rhizobium moroccans* sp. nov.
- Type strain: AGC 32^T^ (=CCMM B1344^T^).
- Accession numbers: 16S rRNA gene, PV739404; whole genome, SRR29855735.
- BioProject: PRJNA1133887; Biosample: SAMN43406629.
- DNA GC content: 61.62%.
- Genome size: 5.24 Mb (Fig 5a).
- ANI/closest relative: 79.9% %.
- dDDH: 20%.

**Figure 5.**
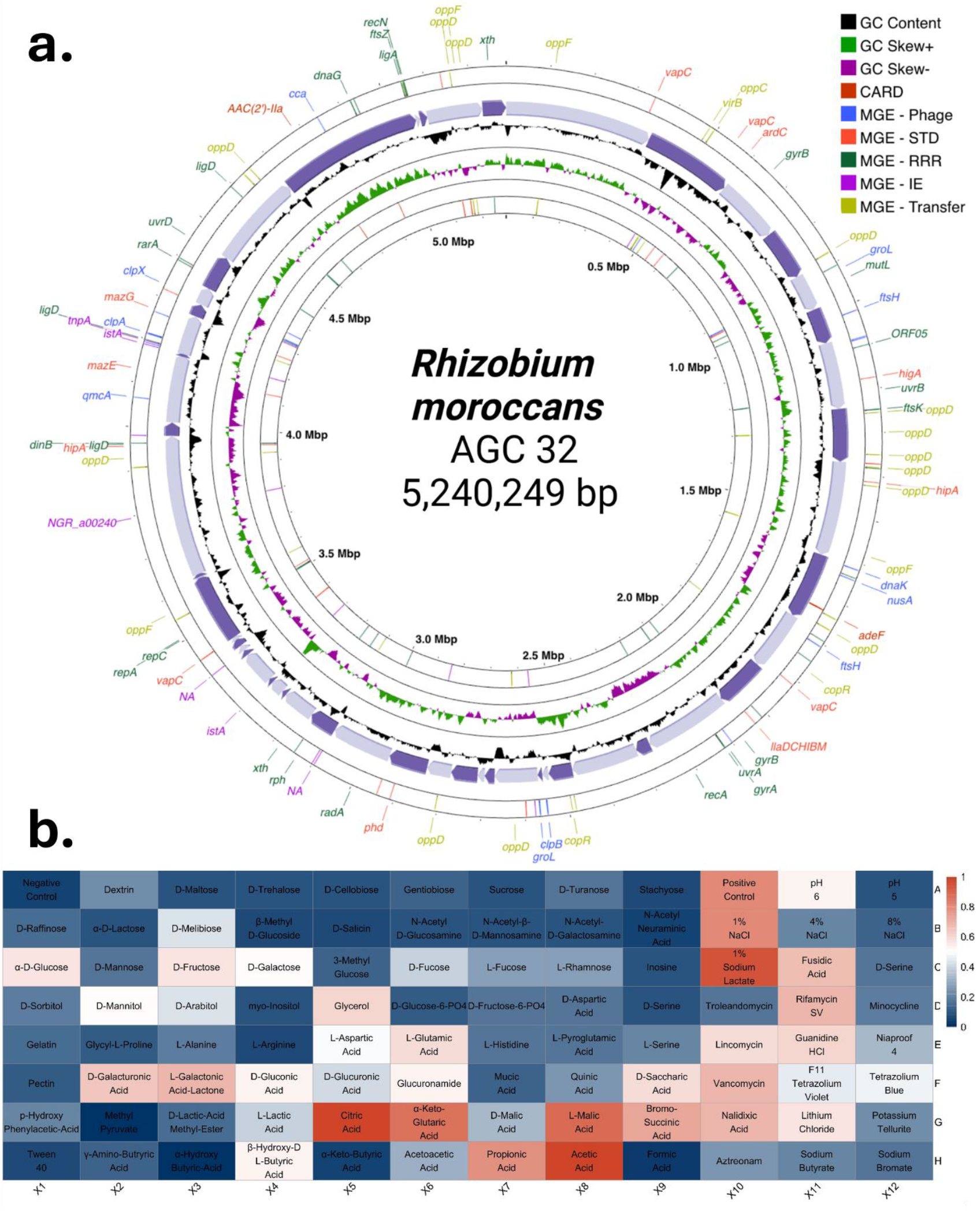
Circular genome map and phenotypic profiling of *R. moroccans* sp. nov. AGC32. (a) Circular genome visualization generated with Proksee, displaying GC skew, predicted genomic islands, and selected functional gene clusters. (b) Biolog GEN III metabolic profile showing carbon source utilization and chemical sensitivity patterns based on positive and negative reactions.

### Phenotypic and plant growth–promoting traits

Following its designation as a novel species, strain AGC32 was phenotypically characterized using the Biolog GEN III MicroPlate system (Fig. 5b). The metabolic profile generated borderline positive reactions, preventing assignment to any species within the GEN III database. For routine cultivation, the strain was resuscitated on yeast extract mannitol (YEM) agar supplemented with 0.0025% Congo red. The isolate was aerobic and oxidative, exhibiting a chemoorganotrophic metabolism with broad utilization of carbohydrates and organic acids typical of rhizosphere-associated *Rhizobium* species.

In the chemical sensitivity panel, AGC32 grew at pH 5–6 and tolerated up to 1% NaCl. It metabolized multiple carbohydrates, including D-glucose, D-mannose, D-fructose, D-maltose, D-trehalose, D-cellobiose, sucrose, and dextrin. Organic acids utilized included citrate, D- and L-malate, propionate, acetate, formate, and α-ketoglutarate. The strain also assimilated several amino acids, including L-alanine, L-arginine, L-aspartate, L-glutamate, and γ-aminobutyric acid, as well as D-galacturonic acid, L-galactonic acid lactone, and glucuronamide.

Sensitivity to rifamycin SV, vancomycin, minocycline, and nalidixic acid was observed. Genome analysis revealed no virulence loci, toxin genes, or pathogenicity islands typically associated with opportunistic plant or human pathogens.

Qualitative plant growth–promoting (PGP) assays demonstrated growth under nitrogen-limited conditions and solubilization of potassium, silicate, phosphate, and zinc, indicated by halo formation and color change on respective media (Fig. 6). These phenotypic traits are consistent with the genomic detection of genes involved in nitrogen metabolism, phosphorus metabolism, potassium homeostasis, and organic acid production (Fig. 3a).

**Figure 6.**
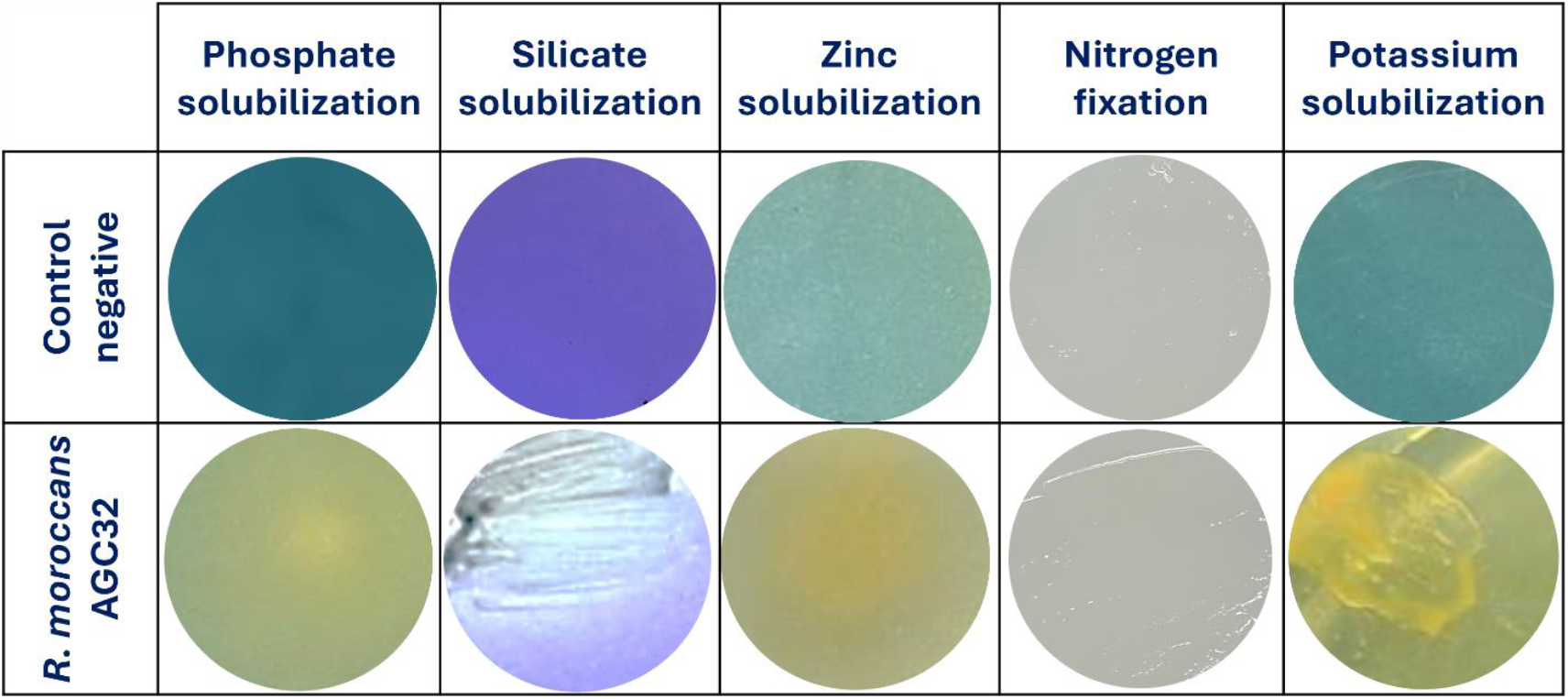
Plant-growth-promoting traits of *R. moroccans* sp. nov. AGC32. Qualitative assays showing solubilization of phosphate, silicate, zinc, and potassium, as well as growth under nitrogen-limited conditions (columns). Rows represent the negative control and strain AGC32. Solubilization of inorganic elements was confirmed by color change in the medium and the formation of visible halo zones around colonies.

## Discussion

Integrative genomic, functional, and phenotypic analyses position *Rhizobium moroccans* sp. nov. AGC32 as a novel endophytic bacterium associated with *Peganum harmala* roots, providing evidence that members of the genus *Rhizobium* can establish non-nodulating, endophytic relationships with non-legume hosts (14, 15). Phylogenomically, AGC32 forms an independent branch within the *Rhizobium* clade, distinct from *R. deserti, R. puerariae*, and *R. leguminosarum*. Genome-based metrics (ANI 79.9%, dDDH 20%, absence of MLST and TYGS matches) confirm species-level divergence, consistent with contemporary prokaryotic taxonomy standards (16). Adapted to arid regions, AGC32’s genomic features provide insights into niche adaptation within the root endosphere of *P. harmala*, highlighting rapid evolutionary diversification among specialist root endophytes.

Beyond taxonomic novelty, AGC32 exhibits a functional genome architecture optimized for survival and plant interaction in arid and semi-arid environments. Its genome encodes genes for protein biosynthesis (132), central carbohydrate metabolism (80), DNA repair (44), oxidative stress response (23), osmotic adaptation (16), and detoxification, reflecting resilience to harsh conditions typical of North African soils, including UV radiation, salinity, and oxidative stress (17). A notable repertoire of flagellar motility genes (47) and membrane transport systems underscores its rhizosphere colonization capacity, essential for establishing plant-microbe interactions (18).

Metabolic reduction, evidenced by limited aromatic compound and amino acid derivative pathways, suggests a specialized strategy adapted to low-complexity nutrient environments, minimizing energetic costs while relying on host-derived metabolites (19). The presence of a complete allantoin utilization operon further supports adaptation to nitrogen-limited root environments (20, 21). The presence of a complete allantoin utilization operon further indicates adaptation to nitrogen-limited root environments, which is characteristic of the natural habitats of *P. harmala* in Morocco, where soils are calcareous, highly mineralized, and generally poor in nutrients, particularly nitrogen (22, 23).

Functionally, AGC32 combines nitrogen fixation (*nifHDK*) with osmoregulated periplasmic glucan transport systems (*yehZYXW*), enabling coordinated adaptation to arid environments with fluctuating nitrogen and osmotic conditions. Additional features, including high-affinity phosphate transporters (*pstSCAB*), catalases (*katG, katE*), TRAP transporters for organic acids, and *nagAB* genes for amino sugar metabolism, reflect overlapping nutrient acquisition and stress-response strategies, with over 20% of coding sequences devoted to nutrient uptake, redox balance, or stress mitigation. This functional redundancy supports enhanced ecological plasticity, consistent with the “gene abundance–niche breadth” hypothesis (24).

Genotype-phenotype integration is supported by Biolog profiling, showing oxidative, strictly respiratory chemoorganotrophic metabolism with extensive carbohydrate, organic acid, and amino acid utilization. Metabolism of citrate, malate, α-ketoglutarate, and γ-aminobutyric acid aligns with root exudate composition, confirming ecological specialization and validating genomic investment in central metabolism.

AGC32 also harbors biosynthetic gene clusters (BGCs) for polyhydroxyalkanoate (PHA), terpene, NRPS-like, RiPP-like, homoserine lactone, and Type I PKS compounds. These clusters suggest multifunctional roles in stress adaptation, carbon storage, antimicrobial activity, quorum sensing, root colonization, pathogen suppression, and modulation of the host environment (25, 26). Low similarity to characterized clusters (MIBiG <0.6) highlights the potential novelty of these secondary metabolites.

Isolation from a non-legume medicinal plant (*P. harmala*) reinforces that rhizobia can act as facultative endophytes beyond classical legume nodulation, contributing to nutrient mobilization, stress tolerance, and plant growth promotion without forming nodules (27). The co-occurrence of nitrogen fixation, mineral solubilization, stress resistance, and specialized metabolic pathways positions AGC32 as a multifunctional endophyte adapted to chemically complex rhizosphere niches, such as the alkaloid-rich root environment of *P. harmala*.

Collectively, these findings highlight distinct ecological strategies of *R. moroccans* and emphasize medicinal plants as reservoirs of metabolically specialized and biotechnologically relevant endophytes. Future work using multi-omics approaches (transcriptomics, metabolomics, proteomics) and chemical characterization of novel BGC products will be essential to validate bioactivity, elucidate plant-growth-promoting mechanisms, and assess potential applications in sustainable agriculture, biocontrol, and drug discovery (28).

## CONSLUSIONS

This study validates our hypothesis that the root endosphere of *Peganum harmala* in Moroccan drylands harbors phylogenetically distinct and genomically adapted *Rhizobium* lineage shaped by both host filtering and arid environmental pressures. In line with our expectations, strain AGC32 represents a novel taxonomic entity within the genus and exhibits genomic features associated with stress tolerance, metabolic versatility, and a plant-associated lifestyle independent of nodulation.

*Rhizobium moroccans* sp. nov., isolated from *P. harmala* roots, is a host-adapted endophyte with a specialized genome supporting plant-growth-promoting traits, stress resilience, and multi-mineral solubilization. Comparative phylogenomics confirms its species-level divergence, while functional analyses reveal genomic signatures consistent with adaptation to arid soils and root-associated niches. The presence of potentially novel biosynthetic gene clusters further highlights the ecological and biotechnological significance of endophyte–host interactions.

Collectively, these findings position *P. harmala* as a valuable reservoir of metabolically specialized rhizobia and expand the ecological framework of the genus *Rhizobium*. Future studies should include controlled inoculation experiments to assess the effects of *R. moroccans* on plant growth, stress tolerance, and symbiotic performance in both legume and non-legume hosts, thereby providing functional validation of its plant-beneficial potential.

## MATERIALS AND METHODS

### Plant habitat and isolation of endophytic bacteria

Healthy individuals of *Peganum harmala* were collected on 27 May 2022 from a natural population in Nzalat Laadam, Benguerir, Morocco (32°06′49.6″N, 7°57′13.0″W) (Fig. S2a). The site is characterized by arid climatic conditions with alkaline soils. Root samples were transported to the laboratory in sterile containers and processed within 24 h (Fig. S2b). Surface sterilization, validation of sterilization efficacy, tissue maceration, serial dilution, plating conditions, colony purification, and long-term preservation procedures were performed as previously described by S. Mouhib et al. (29) (Fig. S3). Briefly, sterilized root tissues were macerated under aseptic conditions and plated on tryptic soy agar (TSA) and potato dextrose agar (PDA). Plates were incubated aerobically at 28°C for 48–72 h, and distinct colonies were purified by repeated streaking (Fig. S3). The isolate designated AGC32 was preserved in 25% (vol/vol) glycerol at −80°C.

### DNA extraction, 16S rRNA gene sequencing, and whole-genome sequencing

Genomic DNA was extracted from overnight cultures grown in TSA broth at 28°C using a protocol of <u>P. Llop et al.</u> (30). DNA quality and concentration were assessed using a NanoDrop spectrophotometer (Thermo Fisher Scientific) and agarose gel electrophoresis.

The 16S rRNA gene was amplified using universal primers 27F and 1492R under standard PCR conditions. PCR products were purified and sequenced by Sanger sequencing. Sequence identity was determined using BLASTn against the NCBI database (29).

Based on 16S rRNA gene affiliation and colony morphology, isolate AGC32 was selected for whole-genome sequencing as described by <u>S. Mouhib et al.</u> (29) (Fig. S4a).

### Genome assembly and taxonomic analysis

Raw reads were quality-checked using FastQC v0.11.9 and trimmed using Trimmomatic v0.39 with default parameters (31). The overall bioinformatics workflow, including preprocessing, quality control, assembly, and taxonomic assignment, is summarized in Figure S4b and follows the pipeline previously described by <u>S. Mouhib et al.</u> (29) for foliar endophytes of the same host plant. The specific tools, parameters, and thresholds applied in the present study are detailed below.

Genome-based taxonomic analysis was performed using the Type (Strain) Genome Server (TYGS) (32) and PubMLST https://pubmlst.org (33). Digital DNA–DNA hybridization (dDDH) values were calculated using the Genome-to-Genome Distance Calculator implemented in TYGS, and average nucleotide identity (ANI) was computed using FastANI v1.33. Species delineation thresholds were applied as described by <u>S. Mouhib et al.</u> (29).

Phylogenomic relationships were inferred using 13 complete genomes of closely related *Rhizobium* type strains, including strain AGC32. An alignment-free phylogenomic analysis was conducted using SANS v2.5 (34). Whole-genome assemblies were compared at the nucleotide level by computing shared and unique k-mers (k = 31). Phylogenetic signal was inferred from weighted k-mer–derived splits without sequence alignment. Only splits compatible with a strict tree topology were retained. Node robustness was assessed by bootstrap resampling of k-mers (1,000 replicates), retaining splits with bootstrap support ≥ 0.75. The resulting strict consensus tree was exported in Newick format. *Mesorhizobium jarvisii* served as the outgroup. The tree was visualized and manually rooted using Geneious Prime v2023.0.3 (Biomatters Ltd., Auckland, New Zealand).

### Functional annotation, genome visualization, and comparative genomics

Genome annotation was performed using Prokka (35). Functional assignments were obtained using KEGG Orthologs via KofamKOALA (36) and subsystems platform (37). Metabolic pathways and gene counts were grouped into major functional categories and visualized as heatmaps.

Genes associated with endophytic adaptation and plant interaction were curated based on published datasets (38, 39), including genes involved in amino acid metabolism, aromatic compound degradation, carbohydrate utilization, energy production, inorganic ion metabolism, metal resistance, lipid cofactors, nucleic acid metabolism, and stress response. Gene counts were normalized using Z-scores and visualized using RAWGraphs 2.0 as a red–blue heatmap.

Secondary metabolite biosynthetic gene clusters (BGCs) were predicted using antiSMASH with default parameters for the detection of NRPS, PKS, and RiPP clusters. The draft genome was visualized as a circular map using Proksee (40), displaying coding sequences, GC content, GC skew, genomic islands, mobile genetic elements, and antimicrobial resistance genes. Cross-referencing of mobile elements and resistance determinants was performed using MobileOG-db (41) and the CARD database (42).

Pairwise whole-genome alignments were conducted using progressiveMauve with default scoring parameters and the “Move contigs” option enabled. High-weight locally collinear blocks were analyzed to assess synteny conservation and genome rearrangements. The draft assembly of AGC32 was aligned with its closest related species, and locally collinear blocks were visualized using the Mauve viewer and exported as PNG images.

### Phenotypic profiling

Phenotypic characterization was performed using the Biolog GEN III MicroPlate system (Biolog Inc., Hayward, CA, USA) according to the manufacturer’s instructions. The system assesses utilization of 71 carbon sources and resistance to 23 chemical stressors through tetrazolium-based colorimetric reactions. Plates were incubated at 28°C for 72 h, and absorbance was measured at 590 nm using the Biolog MicroStation™ reader. Results were interpreted using the GEN III species database.

### Plant-Growth-Promoting (PGP) characterization

For plant growth–promoting assays, strain AGC32 was cultured on TSA for 24 h at 28°C. Phosphate solubilization was assessed using the National Botanical Research Institute’s phosphate (NBRIP) medium (43). Potassium solubilization was evaluated on Aleksandrov medium containing mica as the potassium source (44). Zinc and silicate solubilization were tested on nutrient agar supplemented with ZnO or silicate, respectively, using bromothymol blue or bromocresol purple as pH indicators (45, 46).

Plates were incubated at 28°C for 5–7 days. Mineral solubilization was indicated by halo formation and color change surrounding bacterial growth. Nitrogen assimilation capacity was evaluated by streaking isolates onto nitrogen-deficient combined carbon (CC) medium (Rennie, 1981). Visible growth after 7 days at 28°C was considered positive. All assays were performed in triplicate following the procedures described by <u>S. Mouhib et al.</u> (29).

## ACKNOWLEDGMENTS

This work was funded by the OCP Nutricrops Project AS-85, which is gratefully acknowledged. We thank Bulbul Ahmed, Nabil Radouane, Khaoula Errafii, Imad Khatour, Safaa Machraoui, Issam Meftah Kadmiri, Derly Andrade-Molina, and Salah Eddine Azaroual for their valuable technical support and assistance in bioinformatics. We also gratefully acknowledge the support and computing resources provided by the Toubkal Supercomputer Team at UM6P (Morocco).

## SUPPLEMENTAL MATERIAL

Table S1. Taxonomic identification of isolate AGC32 based on 16S rRNA gene sequencing using the Sanger method.

Table S2. Summary of features of the draft genome assembly of endophytic isolate AGC32 from *Peganum harmala*.

Table S3. Endophytic traits of strain AGC32 isolated from the root endosphere of *Peganum harmala*.

Figure S1. Colony macro-morphology of isolate AGC32 grown on two different culture media.

Figure S2. Sampling site of the host plant located in an arid area along the road between Benguerir and Marrakech.

Figure S3. Graphical representation of the workflow used for the isolation of bacterial endophytes from *P. harmala*.

Figure S4. Experimental workflow and bioinformatic pipelines for whole-genome sequencing of endophytic bacteria isolated from *P. harmala* roots.

## DATA AVAILABILITY

Data Availability Statement: Accession numbers for the 16S rRNA gene sequences are available in GenBank under PV739404. Genome sequence data for the bacterial isolates have been deposited in the Sequence Read Archive (SRA) under SRR29855735. The type strain of the novel species has been deposited in the Moroccan Coordinated Collections of Micro-organisms (CCMM) (https://www.ccmm.ma), with accession numbers B1344^T^ (*Rhizobium moroccans* sp. nov.). The genome assemblies are publicly accessible on Zenodo : (https://zenodo.org) under the following DOIs: https://doi.org/10.5281/zenodo.18664957

